# Full-length direct RNA sequencing reveals extensive remodeling of RNA expression, processing and modification in aging *Caenorhabditis elegans*

**DOI:** 10.1101/2024.06.18.599640

**Authors:** Erin C. Schiksnis, Ian A. Nicastro, Amy E. Pasquinelli

## Abstract

Organismal aging is marked by decline in cellular function and anatomy, ultimately resulting in death. To inform our understanding of the mechanisms underlying this degeneration, we performed standard RNA sequencing and Nanopore direct RNA sequencing over an adult time course in *Caenorhabditis elegans.* Long reads allowed for identification of hundreds of novel isoforms and age-associated differential isoform accumulation, resulting from alternative splicing and terminal exon choice. Genome-wide analysis reveals a decline in RNA processing fidelity and a rise in inosine and pseudouridine editing events in transcripts from older animals. In this first map of pseudouridine modifications for *C. elegans*, we find that they largely reside in coding sequences and that the number of genes with this modification increases with age. Collectively, this analysis discovers transcriptomic signatures associated with age and is a valuable resource to understand the many processes that dictate altered gene expression patterns and post-transcriptional regulation in aging.

## Introduction

Most organisms experience functional decline over time due to accumulation of cellular damage. The process of aging is known to elicit both coordinated and stochastic changes in gene expression, which can either promote decline or aid in maintenance of cellular homeostasis. The nematode, *Caenorhabditis elegans* is a useful animal model for understanding transcriptional changes underlying aging due to its short lifespan and well-annotated genome ^1–3^. *C. elegans* share a high genetic homology to humans and, like humans, experience a functional and anatomical decline in aging ^3–5^.

RNA sequencing (RNA-seq) has been used extensively to characterize the aging transcriptome. Pharmacological suppression of gene expression changes that coincide with aging in *C. elegans* promotes lifespan, highlighting the important contribution of the transcriptome in maintenance of proper cellular function ^6^. Many age-associated gene expression changes are regulated; however, a large proportion arise from loss of transcriptional and post-transcriptional regulation, including changes to transcriptional elongation rate, splicing fidelity, and mRNA surveillance ^7–9^. Loss of other post-transcriptional regulators, including RNA editing enzymes that convert adenosine to inosine in *C. elegans,* also alter lifespan, suggesting a role for RNA modifications in maintenance of a normal lifespan ^10,11^. This hints at potential important roles for other RNA modifying enzymes in maintenance of lifespan, warranting further exploration.

Previous transcriptomic studies in aging have been limited by typical RNA-seq methodologies, which rely on transcript assembly from short cDNA fragments. Oxford Nanopore Technologies Direct RNA sequencing (Nanopore DRS) overcomes these limitations by reading native RNA strands, eliminating the requirement for fragmentation, reverse transcription and PCR amplification, and allowing for sequencing of full-length mRNAs, with no theoretical upper limit to read length ^12^. This method improves transcript annotations, which are difficult to confidently assemble with short reads and has been used to identify novel splice isoforms and 3’UTRs in *C. elegans* at developmental time points ^13,14^. Transcript annotations with full-length support aid in identifying alternative splicing events that may be biologically relevant.

Nanopore DRS also facilitates the identification of mRNA features that require specialized library preparation methods to be detected with standard RNA-seq, including modified RNA nucleotides and poly(A) tail lengths ^15,16^. Modifications like pseudouridine (Ψ) are increasingly being identified in mammalian mRNA, though the difficulty of detecting these modifications has limited our understanding of their functional roles ^17,18^. Leveraging Nanopore DRS to identify such modifications will enhance our understanding of their impact on mRNA genome-wide. Similarly, poly(A) tails have historically required complex experimental and computational methods to sequence ^19,20^. Using these methods, it was determined that short poly(A) tails are a feature of highly expressed, well translated genes in *C. elegans* and other species examined, suggesting that poly(A) tails are subject to co- or post-transcriptional regulation ^21^. Characterizing poly(A) tail lengths across a variety of experimental conditions will further strengthen our understanding of their regulation and impact on mRNA stability and translation.

Despite its many advantages, Nanopore DRS remains limited by its relatively low base-calling accuracy and depth of sequencing, though by both of these metrics the methodology is improving ^22^. Previous studies in *C. elegans* demonstrated a requirement for stringent filtering of raw Nanopore DRS data to identify high confidence mRNA isoforms ^13,14^. Isoform annotations can be further improved using higher accuracy short reads to correct splice junction sequences^23^. Using Nanopore DRS in tandem with short read RNA-seq, therefore, is optimal for performing in depth, comprehensive transcriptomic profiling.

In this study we performed Nanopore DRS in conjunction with short read Illumina RNA-seq across an adult time course in wild-type *C. elegans.* We generated an extensive dataset, which we used to identify novel transcript isoforms and 3’UTRs with full-length support. With these data, we also characterized signatures of the aging transcriptome, including changes to gene expression, poly(A) tail lengths, and an increase in detected inosine and pseudouridine RNA modifications.

## Results

### Long-read and short-read sequencing over an adult time course in *C. elegans*

To better understand how the RNA transcriptome changes during aging in an intact animal, we have profiled RNA expression at eight time points in wild-type (WT) *C. elegans* adults, spanning reproductive (adult days 1-4) and post-reproductive (adult days 5, 7, 10, 15) periods. We chose to profile WT animals, as chemical and genetic manipulations to prevent reproduction alter gene expression and aging pathways ^24,25^. By day 15, about 50% of the population is alive, so we ended collections then to avoid escalation of survivor bias in our gene expression analyses (Figure 1A). Care was also taken to minimize contamination of the adult samples with eggs, progeny, and deceased animals.

**Figure 1.**
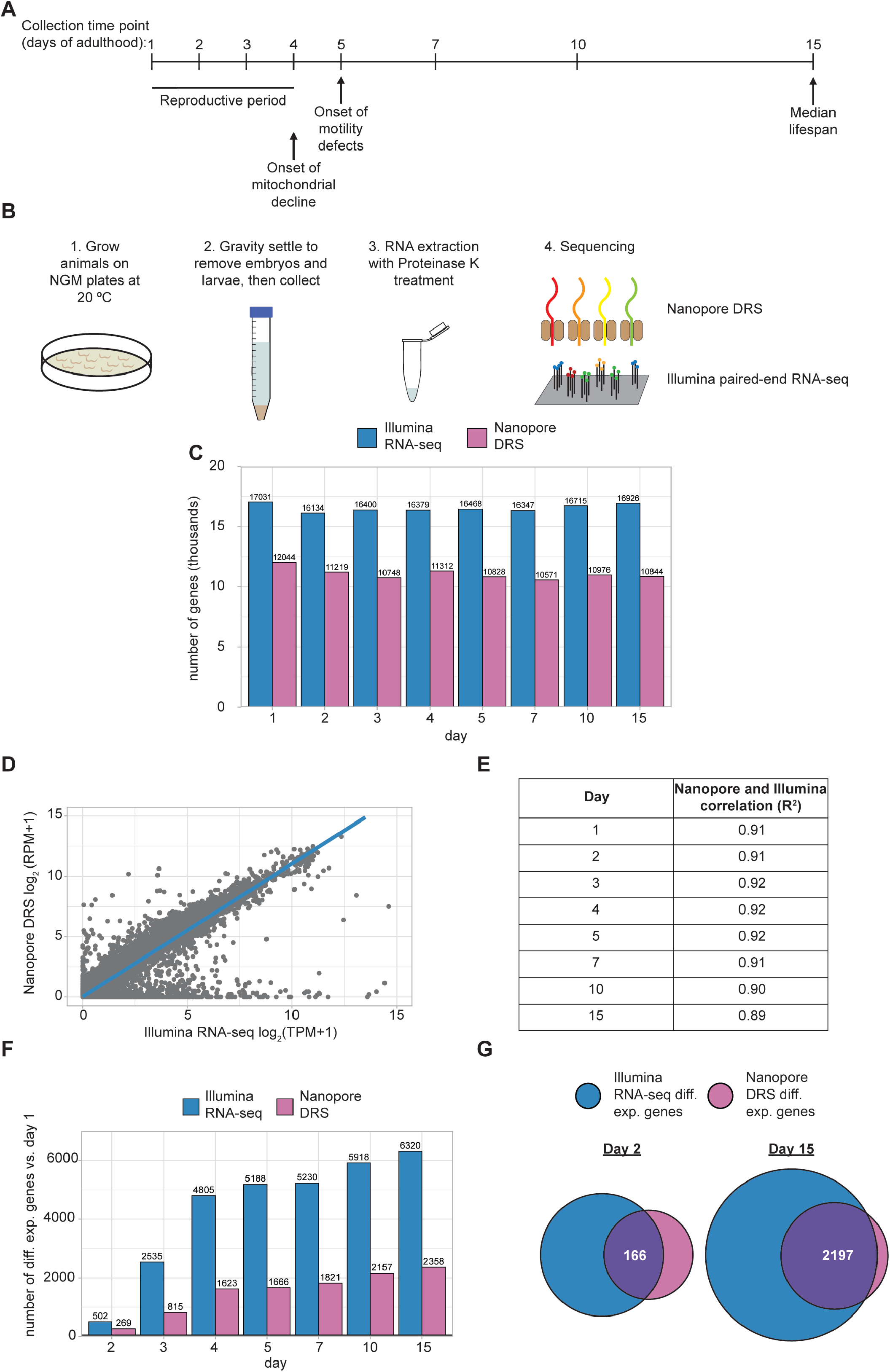
Overview of tandem Nanopore DRS and Illumina paired end RNA-seq. **A.** Collection time points for sequencing experiments. Major physiological changes that occur with aging are indicated in relation to collection time points as reviewed by Spanoudakis and Tavernarakis ^5^.**B.** Overview of sample collection, preparation, and sequencing approach. **C.** Bar plot showing the number of genes detected with Nanopore DRS and Illumina RNA-seq at each time point across three independent biological replicates. **D.** Representative scatterplot comparing the day 1 mean normalized expression of individual genes across three independent biological replicates between Nanopore DRS (log_2_ TPM + 1) and Illumina RNA-seq (log_2_ RPM + 1). **E.** Table showing the R-squared value for mean normalized expression for Nanopore DRS (log_2_ TPM + 1) and Illumina RNA-seq (log_2_ RPM + 1) of individual genes across three independent biological replicates at each time point. **F.** Bar plot showing the number of differentially expressed genes (DESeq2 FDR-corrected p-value ≤0.05) at each time point relative to day 1 for Illumina RNA-seq and Nanopore DRS. **G.** Venn diagrams showing the number of genes that are differentially expressed (DESeq2 FDR-corrected p-value ≤0.05) in Illumina RNA-seq and Nanopore DRS at day 2 and day 15 relative to day 1.

RNA samples from three independent collections for each time point were subjected to Nanopore direct RNA sequencing (Nanopore DRS) to obtain full-length reads and Illumina RNA sequencing (Illumina RNA-seq) (Figure 1B). As expected, more genes were detected with the Illumina RNA-seq method compared to Nanopore DRS (Figure 1C; Table S1; Table S2), as long-read sequencing methods are known to have a relatively lower sequencing depth ^26^. The methods are, however, well correlated when examining normalized gene expression, despite the highly divergent sequencing protocols (Figure 1D; Figure S1). This high correlation between Nanopore DRS and Illumina RNA-seq is observed at each time point sequenced (Figure 1E).

We next examined gene expression changes at each adult time point relative to our earliest time point, day 1. Many genes are significantly differentially expressed at each time point relative to day 1 and the number of differentially expressed genes increases over the aging time course (Figure 1F). Due to limited sequencing depth, fewer differentially expressed genes are detected with Nanopore DRS, but most of these genes overlap with differentially expressed genes detected by Illumina RNA-seq (Figure 1G).

### Identification of hundreds of novel isoforms and 3’UTRs with Nanopore DRS

A key strength of Nanopore DRS is assignment of reads to individual, full-length isoforms, which facilitates identification of novel isoforms. To assign reads to annotated isoforms and identify novel isoforms, we first needed to apply stringent filters to our Nanopore DRS reads (Figure 2A). One limitation of the Nanopore DRS method is read truncation resulting in reads that are not full-length, which are predominantly 3’ biased ^27^. To remove these reads, we used a filtering pipeline to subtract reads that do not correspond to annotated transcription start sites ^14^, then reads lacking poly(A) tails. This filtering pipeline removed ∼50% of reads at each time point (Figure S2A), but over 19 million reads remained to be used for isoform analysis (Figure S2B; Table S3), far exceeding read numbers obtained in earlier studies using Nanopore DRS in *C. elegans* at developmental time points ^13,14^. The filtered reads do not show the 3’ bias observed for reads prior to filtering, demonstrating the utility of these filtering steps in removing reads that are not full-length (Figure 2B).

**Figure 2.**
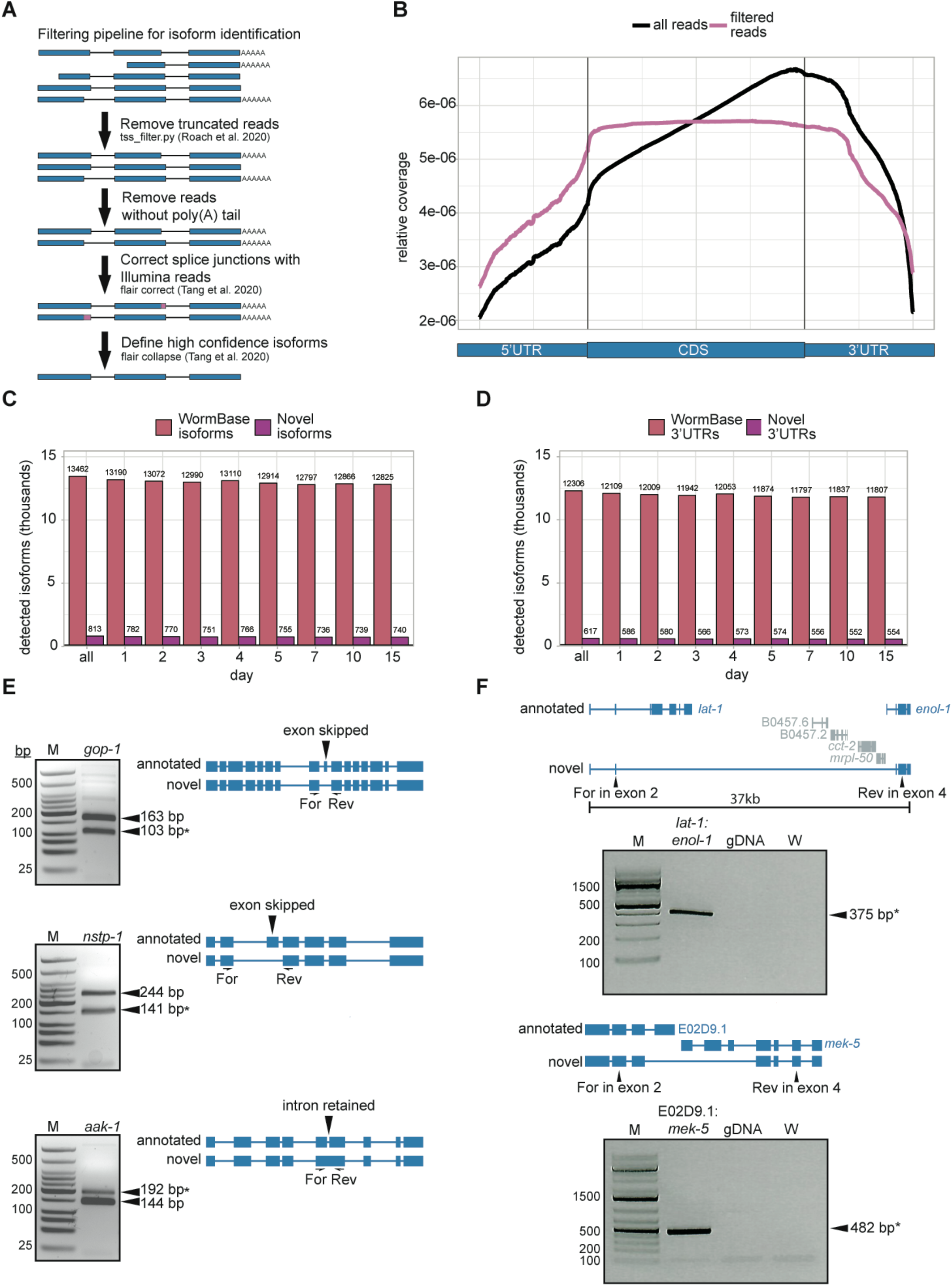
Identification of hundreds of novel isoforms and 3’UTRs. **A.** Approach for filtering and assigning Nanopore DRS reads to define high confidence isoforms. **B.** Metagene plot of normalized Nanopore DRS read density across the average of protein coding genes showing all reads and filtered reads. **C.** Bar plot showing the number of WormBase WS279 isoforms and novel isoforms identified with Nanopore DRS. **D.** Bar plot showing the number of WormBase 3’UTRs and novel 3’UTRs identified using Nanopore DRS. **E.** RT-PCR validation of three novel splicing events for isoforms of *gop-1*, *nstp-1,* and *aak-1*. Agarose gel electrophoresis (*left*) with expected fragment sizes denoted. Asterisk indicates the expected novel isoform product. Schematic representations of the annotated and novel isoforms (*right*) show the novel splicing events with flanking forward and reverse PCR primers used for amplification of cDNA. **F.** RT-PCR validation of two novel isoforms resulting from gene fusions for *lat-1*:*enol-1* and E02D9.1:*mek-5*. Agarose gel electrophoresis (*bottom*) with expected fragment sizes denoted, including genomic DNA and water controls. Schematic representations of the annotated and novel isoform (*top*) show the novel splicing event with forward and reverse PCR primers used for amplification of cDNA.

Filtering steps are necessary for transcript annotation, however they do introduce significant bias, as some genes show a much higher proportion of truncated reads than others. To assess this potential source of bias, we compared filtered Nanopore DRS normalized counts to normalized Illumina RNA-seq counts and found that the methods are no longer highly correlated after filtering (Figure S2C). One example of a gene with an extremely high proportion of truncated reads is *daf-2,* an important regulator of aging ^28^ (Figure S2D). It is not yet clear what causes these truncation events during sequencing, but the observed filtering biases reduce the inherent quantitative nature of the Nanopore DRS approach.

After filtering for full-length reads we used FLAIR to correct Nanopore DRS read splice junctions with higher accuracy Illumina RNA-seq reads, then assigned reads to novel and annotated isoforms ^23^. We applied stringent filters for isoform annotation to eliminate sequences that may represent incompletely spliced transcripts. For this, we required that each isoform represent at least 10% of the total expression for a given gene and set a 20 read minimum for each isoform across all time points. Through this method, we detected over 14,000 isoforms, including 813 novel isoforms representing 782 genes, that are not present in the WS279 WormBase annotation ^29^ (Figure 2C; Table S4). The vast majority of novel isoforms are detected at all adult time points, as low abundance isoforms are likely filtered out by our stringent cutoffs, making it more difficult to detect isoforms only expressed at a single time point. Less stringent filters could be used to pull out more novel isoforms from this dataset, if desired. We also used our isoform annotations identified with FLAIR to extract 3’UTR coordinates and identified 617 novel 3’UTRs in our data, whose 3’ ends do not fall within 10 bp of annotated WS279 WormBase 3’UTR coordinates (Figure 2D; Table S5).

We used stringent filtering criteria and supporting read requirements to provide confidence that our identified novel isoforms represent detectable gene products. We validated new splice isoforms for three genes using RT-PCR with primers flanking novel alternative splicing events. Through this method, we were able to detect PCR products corresponding to annotated as well as novel isoforms for each of the three genes tested, supporting the validity of our detection and annotation method (Figure 2E).

We were also curious to see if any isoforms arise from a fusion of two or more genes, as these isoforms would be difficult to detect with short-read sequencing. Gene fusions are usually aberrant and have known roles in promoting tumorigenesis ^30,31^ and in some cases arise from transcriptional readthrough under stress conditions ^32–34^. Long reads from Nanopore DRS capture full-length transcripts, so are useful in identification of gene fusions, which may span large genomic regions. To screen for this genome-wide, we searched our long-read supported isoforms for introns longer than the median *C. elegans* gene and an overlap with two or more annotated genes. This analysis revealed two interesting fusion isoforms of detectable abundance, both of which we were able to validate using RT-PCR with isoform-specific primers and Sanger sequencing of PCR amplicons (Figure 2F). Interestingly, the fusion isoform spanning the genes *lat-1* and *enol-1* contains a very long (32kb) intron, which overlaps several other genes. This interesting isoform appears to be a novel splicing event rather than an artefact of genomic misannotation, as we do not detect a PCR product with genomic DNA.

To detect more multi-gene isoforms, we deployed LongGF, which searches Nanopore DRS data for reads with multiple genome alignments to identify predicted gene fusions ^35^. This tool identified several gene fusion events across all time points, including inter- and intra-chromosomal gene fusions, which may result from chromosome structural variants or trans-splicing, respectively (Figure S3A). Expression of these fusion isoforms is varied over aging (Figure S3B). Supporting this detection method, we were able to validate the most highly abundant gene fusion isoform, which maps to genes on chromosomes II (C18H9.6) and IV (*clec-173*), by RT-PCR and Sanger sequencing of the resulting amplicon (Figure S3C). Additionally, one of the gene fusion isoforms detected in our analysis, *eri-*6:*eri-*7, was previously reported to arise from trans-splicing of two separate pre-mRNAs ^36^.

Detection of these rare gene fusion isoforms, novel splice isoforms, and novel 3’UTRs highlights a distinct advantage of the Nanopore DRS method. Our use of adult time points and our high depth of sequencing build upon previous Nanopore DRS studies in *C. elegans* and will serve to strengthen genome annotations for the community.

### Accumulation of distinct transcript isoforms during aging

Isoforms with full-length support help validate and update existing gene annotations, but the low read numbers and bias introduced from filtering Nanopore DRS reads make it difficult to examine differential isoform usage at a large scale with these data. Therefore, we leveraged our Illumina RNA-seq data to perform alternative splicing analysis with SUPPA2 ^37^ at each of our time points relative to day 1, using isoform annotations with full-length Nanopore DRS support to define exon boundaries. We detected several alternative splicing events at later days relative to day 1, including in genes required for stress resistance and proper locomotion (Figure 3A; Table S6). These alternative splicing events include exon skipping, use of mutually exclusive exons, intron retention, and alternative first exon usage (Figure 3B). They may be important for regulation of expression of these genes or protein function over the course of aging.

**Figure 3.**
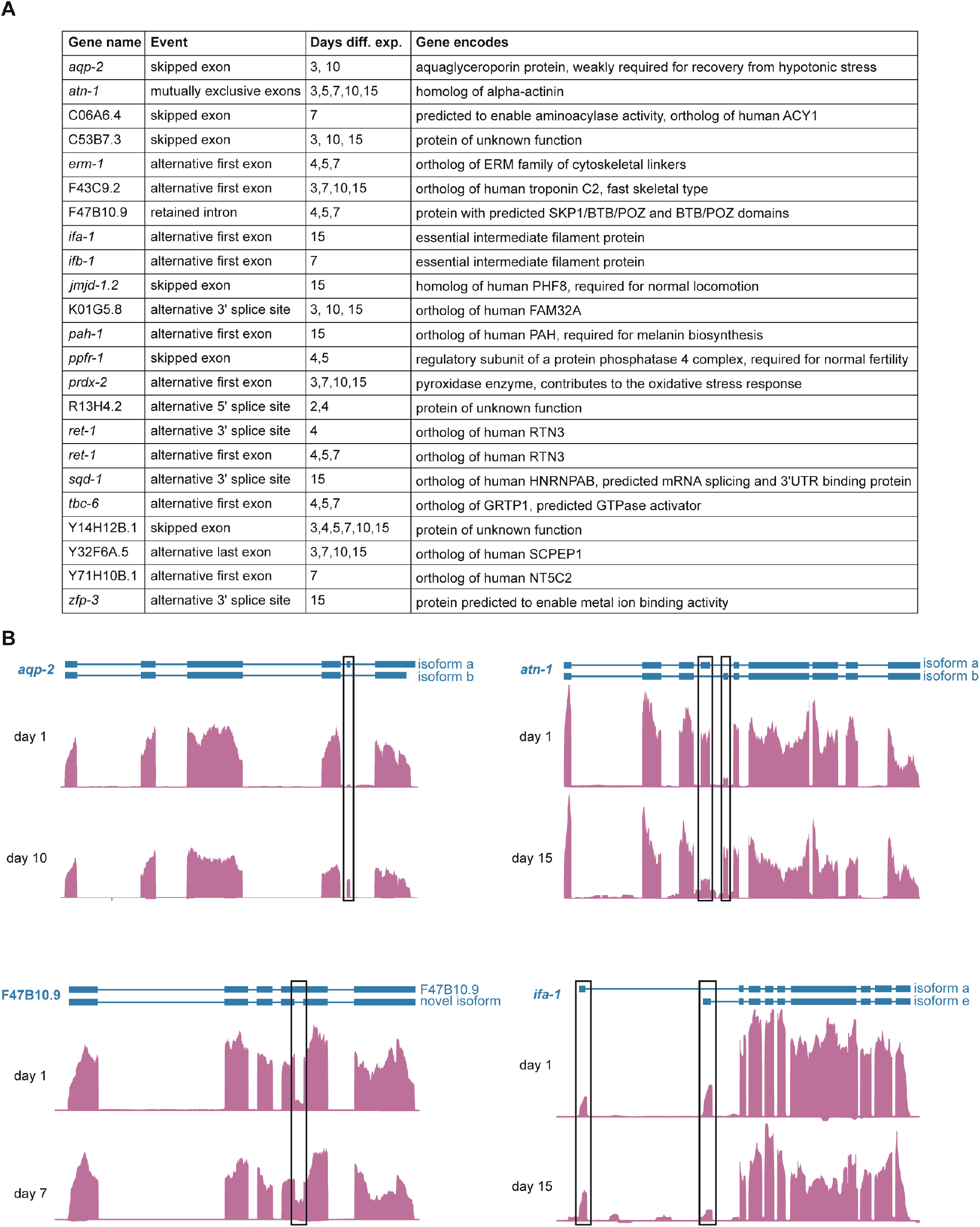
Alternative transcript isoforms in aging. **A.** Table showing all genes with significant (*P*<0.05) change in relative abundance (PSI) of an alternative exon inclusion event relative to day 1 with brief descriptions of known or predicted gene functions from WormBase. **B.** Genome browser screenshots of representative genes (*erm-1, atn-1,* F47B10.9, and *ifa-*1) with alternative exon inclusion events at day 1 and denoted adult time points. Exons with differential PSI at the denoted time point are highlighted with black boxes.

### Global changes in RNA processing and poly(A) tail lengths during aging

To forge a deeper understanding of features of mRNA processing that may change with aging, we examined global trends in young animals compared to old animals. We combined our three earliest time points (days 1, 2, and 3) and our three latest time points (days 7, 10, and 15) to compare young and old animals, respectively. To get a better idea of how splicing and transcription may be altered in aging, we sought to examine the percent of reads in introns and intergenic regions, as well as the use of unannotated splice junctions. For these analyses, we used Illumina RNA-seq reads due to their higher accuracy and read numbers.

We reasoned that changes to the percent of total reads mapping to introns may indicate splicing defects, altered splicing efficiency, or a change in transcription rate. When we strictly defined introns, requiring they not overlap with any WormBase WS279 annotated exon sequence, we did not see evidence of increased intron inclusion in old animals compared to those in young, as the percent reads mapping to introns is not significantly different (Figure 4A; Table S7). This result is contrary to a previous report where a slight increase in intronic read assignments during aging was observed ^8^. This disparity may result from our more stringent definition of introns or the use of combined multiple young and old time points. A similar analysis examining the percent of reads assigned to intergenic regions, which can serve as a proxy for inefficient transcription termination or improper transcription start site selection, revealed a modest but significant increase in intergenic region assignment in aging (Figure 4B).

**Figure 4.**
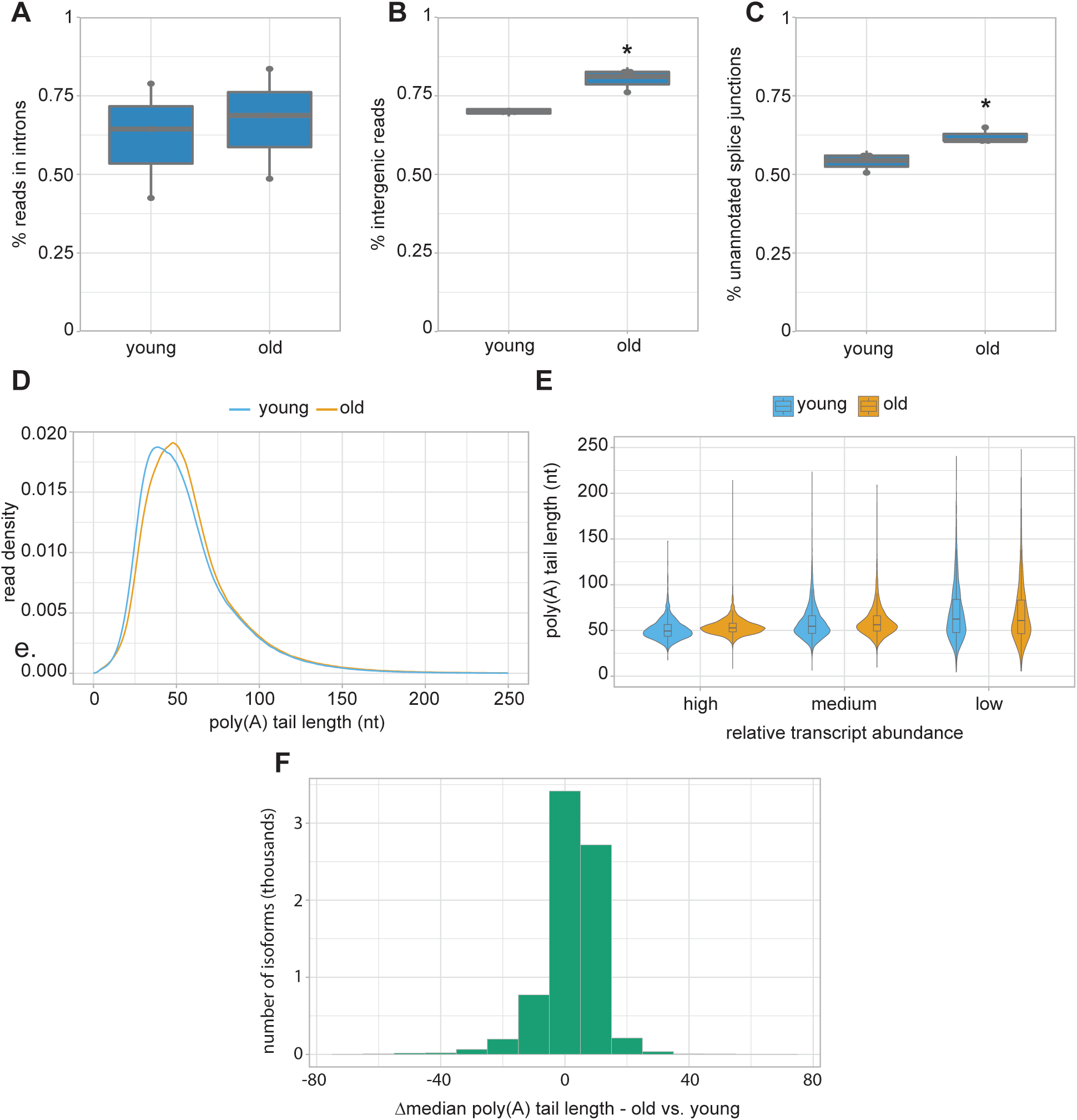
Altered RNA processing in aging. **A.** Boxplot (thick horizontal bar represents the mean, thin horizontal bars represent the interquartile range, and the vertical line represents the 95% confidence interval, dots represent data for individual replicates) showing the percent of Illumina RNA-seq reads mapping to introns. *P*= 0.7417 (*t-*test, two-sided after probit transformation). **B.** Boxplot showing the percent of Illumina RNA-seq reads mapping to intergenic regions. \**P*= 0.04402 (*t-*test, two-sided after probit transformation). **C.** Boxplot showing the percent of splice junction reads that do not match WS279 annotated splice junctions. \**P*=0.029 (*t-*test, two-sided after probit transformation). **D.** Density plot showing poly(A) tail length distributions. *P<*2.2e-16 (Kolmogorov-Smirnov test). **E.** Violin plot with inlaid boxplot of median poly(A) tail length distribution for isoforms grouped by relative abundance (low = 1st quartile, medium = 2nd and 3rd quartiles, high = 4th quartile, based on normalized isoform expression). *P<*2.2e-16 (Kolmogorov-Smirnov test) between expression groups in young and old animals. **F.** Histogram of change in median isoform poly(A) tail lengths in old compared to young animals.

To examine splicing fidelity, we looked at the percent of splice junction reads that do not match annotated splice junctions, which was previously reported to increase in day 15 adults ^8^. Here, we also observe a slight but significant increase in the percent of reads that do not match WormBase WS279 annotated splice junctions (Figure 4C), suggesting a modest accumulation of aberrant transcripts in old animals compared to young. We observe no bias for change in 3’ or 5’ mismatches for unannotated splice junction reads (Figure S4), which may indicate that this accumulation is due to a reduction in their targeted degradation, or that both 5’ and 3’ splice site recognition is affected in later adulthood.

Another feature of mRNA processing that we hypothesized may be altered in aging is polyadenylation. Due to limitations of typical sequencing methods, poly(A) tail lengths have not yet been examined in this context. Here, we again compared young and old animals, first looking at bulk poly(A) tail length distributions from our Nanopore DRS data. We find that poly(A) tails display a modest increase in size in old animals (median=53 nt) compared to young (median=50 nt) (Figure 4D). We note that adult poly(A) tail length distributions and median tail lengths closely resemble those of larval stage *C. elegans* ^13,14,21^.

There is a known anticorrelation between transcript abundance and poly(A) tail length in larval *C. elegans* and other species ^21^. To determine if more highly expressed transcripts maintain shorter poly(A) tails in adults, we binned transcripts by abundance and examined median poly(A) tail length distributions in these groups. This analysis reveals that despite the overall tail lengthening observed in older animals, highly abundant transcripts maintain shorter poly(A) tails than less abundant RNAs (Figure 4E). This implies that poly(A) tail length is likely subject to the same regulatory processes in aging as in larval stages.

To determine if any specific transcripts undergo large changes to median poly(A) tail length over aging, we compared the change in tail lengths for individual isoforms in old animals compared to young (Figure 4F; Table S8). This analysis revealed that the vast majority (81%) of isoforms undergo very little change to poly(A) tail lengths, with a change in median poly(A) of less than 10 nt in old animals compared to young. Only 3.8% of isoforms undergo large changes to median poly(A) tail lengths of more than 20 nt. Most large changes to poly(A) tail lengths can be explained by changes to transcript abundance, in agreement with the previous observations. However, some genes are notable exceptions, with large age-dependent changes to poly(A) tail lengths without corresponding changes in abundance. This subset includes genes in stress response, fatty acid binding, and heme transport pathways (Figure S5). Together, these results show there are distinctive global changes to post-transcriptional RNA processing, including decreased splice site fidelity, increased intergenic RNA expression, and poly(A) tail lengthening, that coincide with aging. However, these important regulatory programs appear to be largely maintained into late adulthood despite the immense changes to physiology and gene expression programs that occur in aging.

### Abundance and frequency of inosine edits increase in aging

Finally, we leveraged our Nanopore DRS and Illumina RNA-seq data to identify modified RNA nucleotides and determine if their abundance and distribution change in aging. We specifically searched for signatures associated with adenosine to inosine (A-to-I) editing and pseudouridine (Ψ). A-to-I editing can be easily detected with Illumina RNA-seq data, as inosine is reverse transcribed as guanosine during library preparation ^38^. So, we queried for adenosine to guanosine (A-to-G) mismatches in our Illumina RNA-seq with SAILOR ^39^, to take advantage of the method’s high depth of sequencing. We combined young (days 1, 2, 3) and old (days 7, 10, 15) time points and required high confidence edits present in more than 5% of reads covering each site and in all three biological replicates.

With these stringent filters applied, we identified over 3,800 A-to-G edits in both young and old animals, which we consider putative A-to-I editing sites (Table S9). The majority of these sites lie in 3’UTRs (Figure 5A), consistent with previous reports in human and *C. elegans* ^40–42^.

**Figure 5.**
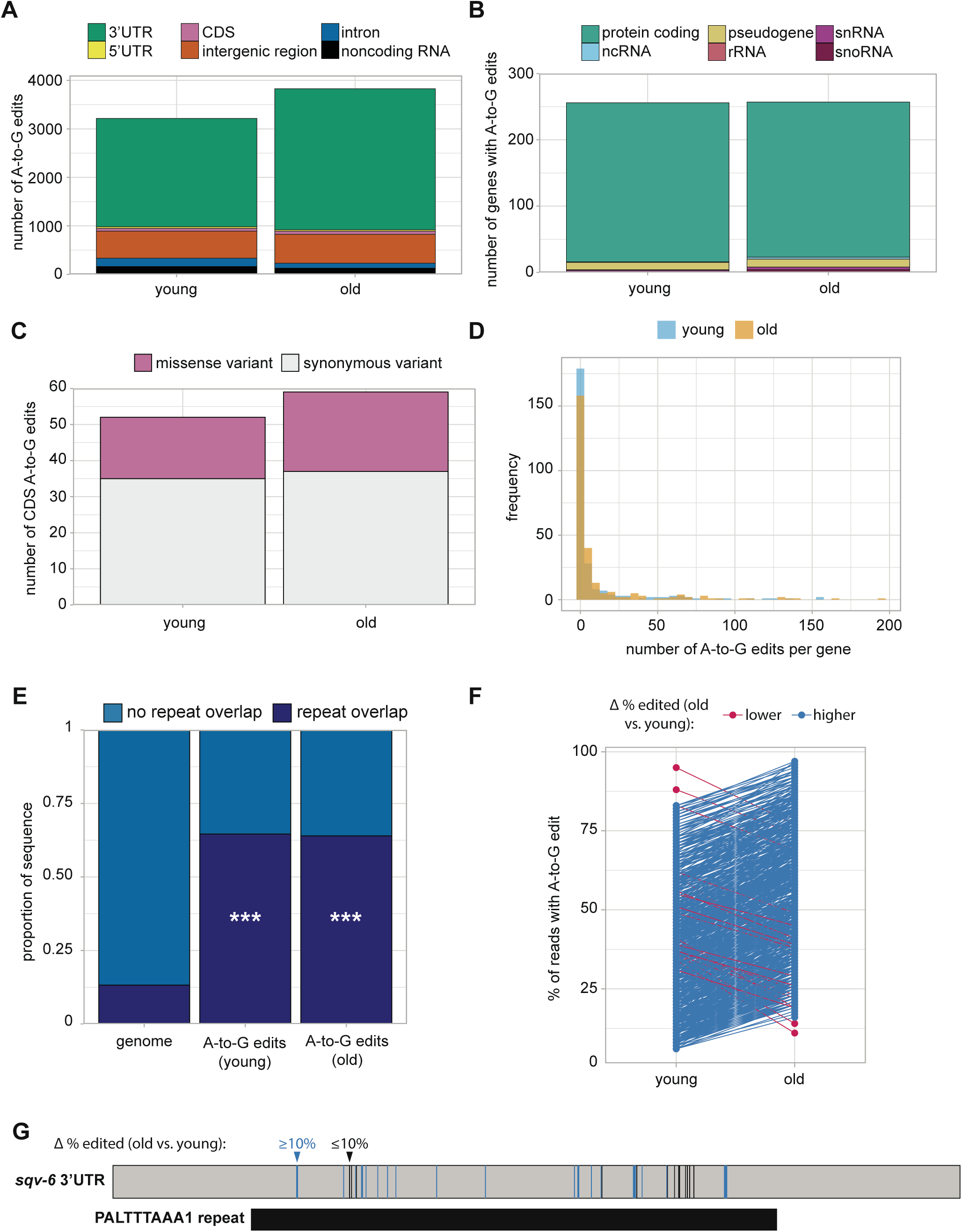
Genome-wide detection of inosine editing. **A.** Stacked bar plot showing the number of A-to-G edits that map to each gene feature. **B.** Stacked bar plot showing the biotype of genes with A-to-G edit(s). **C.** Stacked bar plot showing the number of A-to-G edits in coding regions predicted to result in missense or synonymous codon variants. **D.** Histogram of the number of A-to-G edits per gene. **E.** Stacked bar plot showing the proportion of A-to-G edits that overlap with repeat sequences and the proportion of repeat sequences in the whole genome. \*\*\**P*<2.2e-16 for young and old compared to the whole genome. **F.** Plot with the percent of edited reads for A-to-G sites with large changes (≥10% lower or higher) in editing in young compared to old animals. Each point represents a single A-to-G edit site. **G.** Illustration of A-to-G edit sites in the 3’UTR of *sqv-6*. Vertical bars represent A-to-G edit sites, colored by change to percent edited reads in old compared to young. All sites in the *sqv-6* 3’UTR overlap with the PALTTTAAA1 repeat.

Though A-to-G sites are detected in a variety of classes of RNA, most sites lie within protein coding genes (Figure 5B). Most genes with inosine have edits in both young and old animals (Figure S6A), but there are more inosine sites detected in old animals (Figure 5A), suggesting there may be age-dependent regulation of inosine editing.

Inosine in mRNA coding regions has the capacity to alter translation through recoding or translational stalling ^43^. We wanted to assess whether detected inosine edits in coding regions could change the amino acid encoded at each position. Inosine is most often interpreted as guanosine by translation machinery ^43,44^, so we asked whether substitution of guanosine at edited sites would alter the encoded amino acid. Many inosines indeed give rise to missense variants (Figure 5C), with the potential to alter protein function. Among the predicted missense variants resulting from inosine editing, the majority at both time points change the chemical properties of the encoded amino acid (Figure S6B). Therefore, while less abundant than their 3’UTR counterparts, many inosine edits in coding sequence may impact protein function.

Many genes contain multiple inosine edits in both old and young animals (Figure 5D), consistent with previous reports in *C. elegans,* where clusters of inosine edits in 3’UTRs overlapping with repetitive sequence elements were observed ^45^. We also detect a significant enrichment for overlap of inosine edits with repeat sequences at both young and old time points, when compared to the abundance of repetitive elements genome-wide (Figure 5E).

Furthermore, over 20% of the inosine edits located in 3’UTRs overlap potential microRNA binding sites (Table S9). As inosine has different base-pairing properties than adenosine ^46^, these changes may alter the efficiency of microRNA targeting and regulation of these genes.

Because we observed an overall increase in the number of high confidence inosine edits in aging, we also asked if individual sites are more frequently edited in old animals. To this end, we looked at sites that are detected in young and old animals and compared the percentage of reads with A-to-G edits at each time point. Of the 2,659 inosine sites detected at both young and old time points, 904 inosine sites have more than a 10% change in editing in old compared to young animals. Of these, almost all (98%) increase in editing percentage (Figure 5F). This result suggests that there is an age-dependent increase in inosine editing genome-wide in *C. elegans.* This global trend is also apparent for individual genes with multiple inosine edits (Figure 5G). Overall, our results reveal an increase in the total number of adenosine to inosine conversions in aging, as well as a striking increase in the percentage of edited transcripts at many sites.

### Evidence for pseudouridine editing in *C. elegans*

Pseudouridine is not detectable with Illumina RNA-seq without special methods for library preparation ^47^, and its presence has not yet been examined genome-wide in *C. elegans*. We used a newly developed program, NanoPsu ^48^, to detect Ψ in our Nanopore DRS sequencing data, as it does not require specialized sample preparation. To ensure we only consider high confidence Ψ sites, we required that sites have a ≥0.90 confidence score and be detected in all three biological replicates. With these criteria, we detect 312 Ψ sites in young and 476 sites in old animals (Table S10). Most of these sites lie within coding regions, where they may potentially affect translation efficiency or fidelity (Figure 6A). Almost all detected Ψ sites are within protein coding genes (Figure 6B), but we also see Ψ sites in ribosomal RNA (rRNA) genes, which we recover to some extent despite the Nanopore DRS selection for RNAs with poly(A) tails due to their extremely high abundance. rRNAs are known to be modified with Ψ in many species ^49^, so their presence in this analysis strengthens our confidence in the detection method.

**Figure 6.**
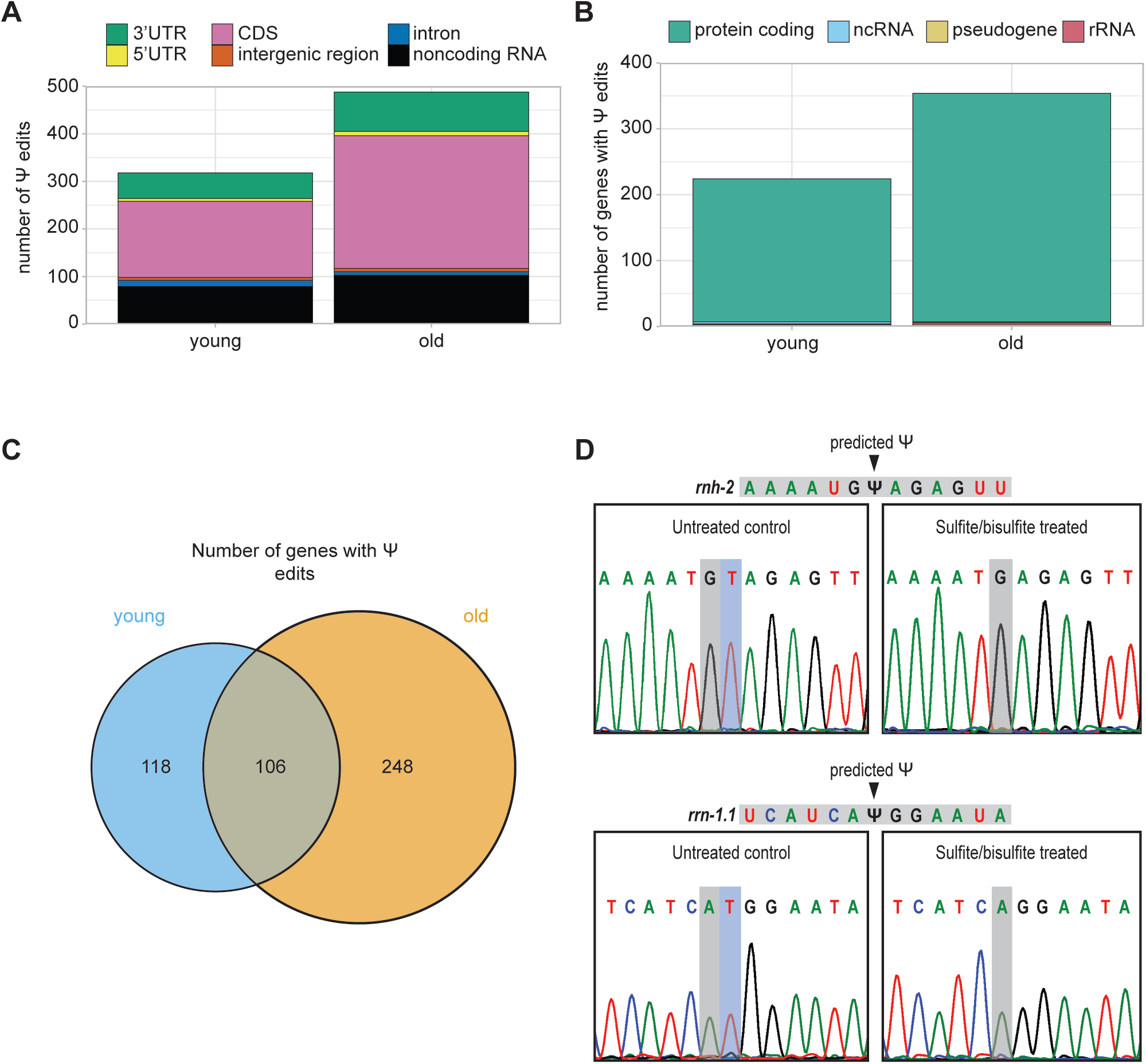
An age-dependent increase in pseudouridine editing. **A.** Stacked bar plot showing the number of pseudouridine edits that map to each gene feature in young (days 1, 2, 3) and old (days 7, 10, 15) animals. **B.** Stacked bar plot showing the biotype of genes with pseudouridine edit(s) in young and old animals. **C.** Venn diagram with the number of genes with pseudouridine edits in young and old animals. **D.** Sanger sequencing results from cloned *rnh-2* and *rrn-1.1* PCR products amplified from reverse transcribed sulfite/bisulfite treated and untreated RNA. Predicted pseudouridine sites and resulting deletions are indicated.

Most genes with Ψ only have a single edited site, based on our strict criteria. Less than 5% of these genes have multiple sites and there is no significant motif enrichment for sequences surrounding Ψ sites. Ψ sites in *C. elegans* small nuclear RNAs (snRNAs) have been mapped ^50^, but these genes lack poly(A) tails, so are not well detected with Nanopore DRS and thus do not emerge in this analysis. Around half of the genes with Ψ in young animals also have Ψ in old animals. Strikingly, however, half of detected Ψ genes are unique to the older time points (Figure 6C). Higher detection of Ψ in aging cannot be explained by increased expression of these genes with edits only detected in the older samples, as they have varied expression patterns over aging (Figure S7A).

While this is the first study to identify Ψ in *C. elegans* mRNA, the *C. elegans* genome has numerous predicted pseudouridine synthase genes that are homologous to human, including enzymes that edit mRNA in human ^51^ (Figure S7B). These genes have dynamic expression patterns in aging (Figure S7C). Our detection of Ψ genome-wide and evidence for pseudouridine synthase enzymes in the *C. elegans* genome led us to validate Ψ sites supported by Nanopore DRS. For this, we used a Sanger sequencing-based approach wherein treatment of RNA with sulfite and bisulfite introduces a deletion at Ψ sites during reverse transcription ^52^. With this approach, we detected a deletion at predicted Ψ sites in the coding region of the *rnh-2* mRNA and in the rRNA *rrn-1.1* upon sulfite/bisulfite treatment (Figure 6D). This validation provides strong evidence of Ψ in *C. elegans* mRNA.

### New features of known aging genes

To summarize, we identified new characteristics that define the aging transcriptome, including a global increase in inosine and pseudouridine modifications and a small, but significant, decline in RNA processing fidelity. We also identified novel splice isoforms and 3’UTRs and instances of age-dependent accumulation of specific isoforms. Particularly interested in annotating new features of genes with known roles in aging, we compared our findings to a list of 1,484 genes previously reported to promote or antagonize longevity ^29,53,54^. We find that many newly uncovered features identified in this study do indeed overlap with known regulators of aging (Figure 7; Table S11). In all, we annotated previously unknown features of 227 genes with published roles in aging. Identification of such features may be important for understanding how these important genes are regulated. For example, *aps-3,* which is involved in intracellular membrane trafficking and antagonizes longevity ^29^, has over 100 inosine edits and a pseudouridine site within its 3’UTR. Similarly, some genes that are alternatively spliced are known regulators of lifespan, including *erm-1* and *prdx-2* ^55,56^, which both use alternative first exons in aging relative to day 1. We identified and validated a novel gene fusion isoform of *enol-1*, which promotes longevity in *C. elegans*. Human orthologs of this gene are implicated in age-related diseases including Alzheimer’s disease and cancer ^57^. These examples highlight some of the new information learned about important aging regulators through this study.

**Figure 7.**
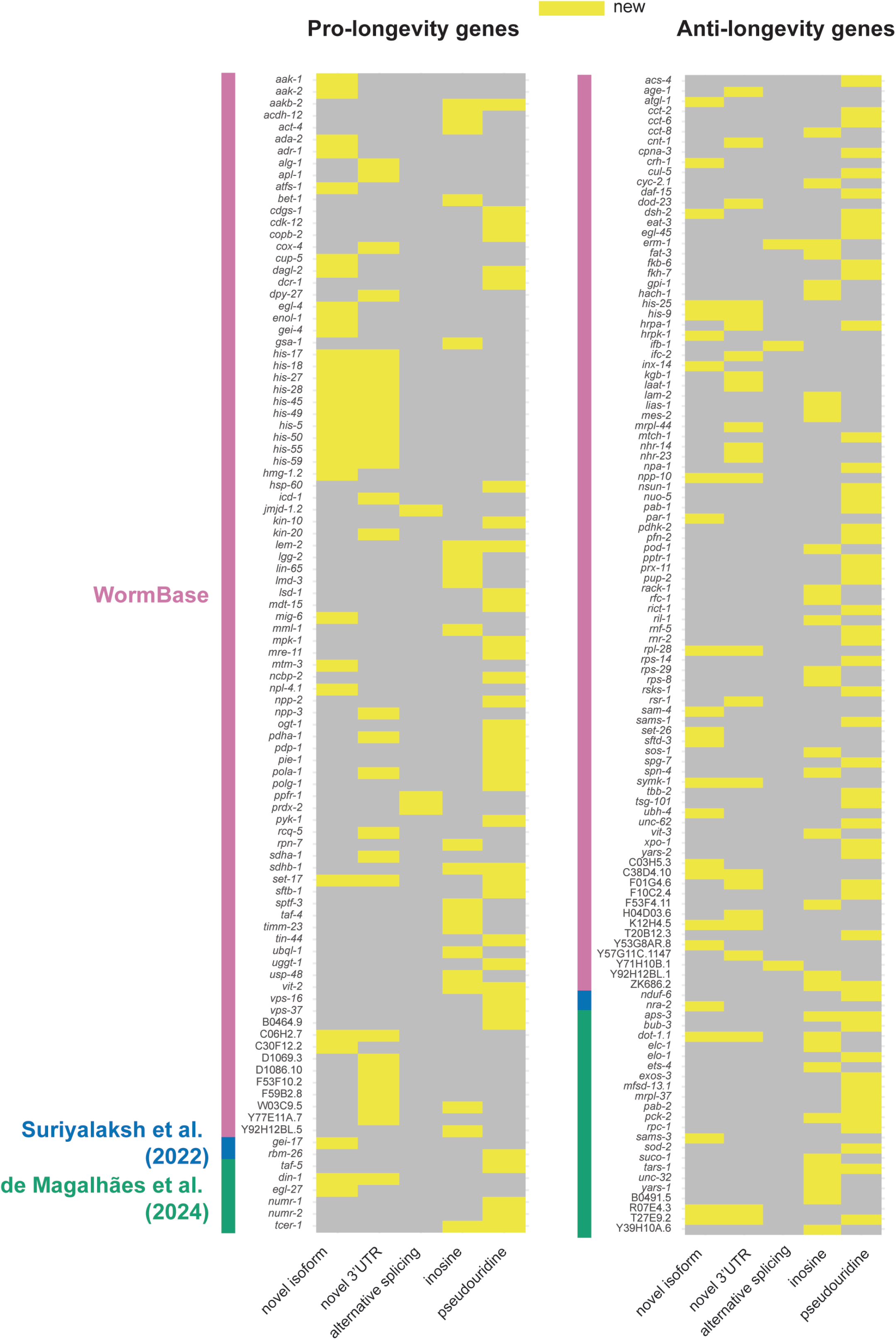
New features of hundreds of known aging genes. Heatmaps of pro- or anti-longevity genes with new features (novel isoforms, novel 3’UTRs, age-dependent alternative splicing, inosine, pseudouridine) identified in this study. An initial list of 1,484 aging genes was generated from RNAi or loss of function (LOF) allele phenotypes from WormBase (anti-longevity=extended lifespan, pro-longevity=reduced lifespan) ^29^ and additional non-overlapping annotated lifespan regulating genes from the literature ^53,54^. Colored side bars on the heatmaps indicate the annotation supporting each gene.

## Discussion

In this study we performed an in-depth investigation of transcriptome changes that occur over an aging time course in wild type *C. elegans*, which we grew under standard laboratory conditions. We generated a comprehensive dataset using the well-established Illumina RNA sequencing method in tandem with long-read Oxford Nanopore Technologies direct RNA sequencing. With Nanopore DRS we generated over 19 million full-length reads, allowing for the identification of 813 novel isoforms, including novel gene fusion isoforms, and 617 novel 3’UTRs, which will improve existing gene annotations and serve as a resource for *C. elegans* researchers. By comparing older to younger populations to identify signatures of aging, we observed age-dependent alternative splicing and measurable changes to RNA processing, including an increase in unannotated splice junctions and reads in intergenic regions. Nanopore DRS reads allowed us to assay poly(A) tail lengths genome-wide, where overall we observed adult poly(A) tails similar in length to those in larval stages, with a modest tail lengthening in aging. The most striking changes coinciding with aging were uncovered when we investigated RNA modifications. We detected an increase in adenosine to inosine editing, both in the number of genes with these edits and the editing frequency at over one third of sites. Using Nanopore DRS to detect pseudouridine sites genome-wide, we present the first map of Ψ sites in *C. elegans* mRNA. Like with inosine, we observe an age-dependent increase in Ψ sites. With our comprehensive dataset and analyses, we identified novel isoforms, splicing events, and modifications for hundreds of pro- and anti-longevity genes. These findings reveal new features of the aging transcriptome and will serve as a valuable resource for other scientists interested in the transcriptional and post-transcriptional effectors of aging.

### Advantages of tandem long-read and short-read sequencing

In our study, we combined long-read sequencing data with short-read sequencing. This allowed us to answer more biological questions and strengthened our novel isoform annotations. While short-read RNA sequencing has been applied in *C. elegans* and other organisms to assess age-dependent changes to gene expression, these methods were limited by read length and their requirement for reverse transcription and PCR amplification of libraries. We generated reads exceeding 10kb and leveraged the ability of this method to sequence RNA directly to investigate poly(A) tail lengths and pseudouridine modifications genome-wide. Despite major improvements to Nanopore DRS in recent years, reads generated through this method remain lower in accuracy as compared to short reads ^58,59^. Correcting splice junctions of long reads with supporting short reads, as we did with our Nanopore DRS data, accounts for this and allows for higher confidence isoform annotations ^23^. Similarly, filtering for full-length reads was necessary for isoform assignment, as we observed a large proportion of reads that are not full-length. Here, we relied on current genome annotations to define full-length reads, as was previously established ^14^. The 5’ read truncations in Nanopore DRS remain among the biggest limitations of this technology. Specific genes are more susceptible to these truncations, likely due to difficult to read sequences or structures. This biased our quantification of full-length isoforms, which led us to use our short-read data to investigate alternative splicing, further demonstrating the utility of applying both sequencing methods in tandem.

### Alterations to RNA processing in aging are detectable but not widespread

Our adult time course, with collection time points ranging from young adult to day 15, allowed us to ask how RNA processing changes during aging. We identified several genes with alternative splicing in later days relative to day 1, including *ret-1,* which was previously shown to be alternatively spliced in aging ^8^. While only a small proportion of expressed genes were found to be alternatively spliced later in adulthood, alternative isoform usage for individual genes can be highly relevant for longevity. Isoforms generated through alternative splicing, alternative transcription start sites, or alternative polyadenylation can regulate longevity differentially. For example, loss of function alleles of *daf-2* famously result in an extended lifespan in *C. elegans* ^28^, whereas overexpression of a non-signaling insulin receptor isoform, *daf-2b,* extends lifespan ^60^. This example highlights the contribution of alternative splicing in maintenance of a normal lifespan and provides justification for further investigation of the alternative splicing events identified in this study. While detectable shifts in alternative splicing were not widespread, individual alternatively spliced isoforms can play important cellular roles.

To look at global splicing changes we assayed reads in introns and unannotated splice junction reads as in Heintz et al ^8^. Our results agree with their finding that unannotated splice junction use modestly increases with age. We do not, however, see the same increased intron retention they reported in our samples from pooled young and old time points. We believe this is due to our stringent definition of introns, which removed any annotated intron sequence overlapping with an exon of another transcript. Regardless, the effects on splicing we do observe in aging are modest, so it remains unclear whether these small changes are biologically relevant under normal growth conditions. Loss of the key splicing factor SFA-1 does not affect lifespan when animals are fed *ad libitum* but is required for lifespan extension under dietary restriction ^8^. Further, the increase in unannotated splice junction reads does not necessarily indicate a decrease in splicing fidelity in aging. Accumulation of improperly spliced mRNAs could result from a decline in recognition or degradation of aberrant transcripts. For example, nonsense mediated decay (NMD) activity declines with age in *C. elegans* ^61^, which could explain these observations.

We also observed a modest overall change to poly(A) tail lengths in aging. Poly(A) tail lengths are anticorrelated to transcript abundance and codon optimality ^21^, hinting at a link between regulation of poly(A) tail length and translation. We do observe tail lengthening in aging, but fluctuations in tail length resemble similar fluctuations observed between larval stages in *C. elegans* and, thus, likely do not represent meaningful deterioration of tail length regulation ^13,14^. Some mRNAs experience larger fluctuations to poly(A) tail lengths over our time course and, thus, may be interesting candidates for further investigation of poly(A) tail length dynamics in an intact animal.

### Inosine editing increases in an age-dependent manner

We observed a striking increase in the number and frequency of inosine edits during aging in this study. This is particularly interesting because loss of function alleles of the *C. elegans* adenosine deaminase acting on RNA (ADAR) enzymes, which catalyze conversion of adenosine to inosine, alter lifespan ^10,11,62^. Specifically, animals lacking ADR-2, which actively deaminates RNA, are long-lived, and animals lacking ADR-1, which regulates editing efficiency through interactions with ADR-2 and RNA, are short-lived ^10^. Based on the *adr-2* loss of function phenotype and the increase in inosine editing that correlates with age, it may stand to reason that increased inosine edits in older animals promote degenerative aging phenotypes. Inosine editing has also been linked to longevity in humans, where single nucleotide polymorphisms (SNPs) in ADAR enzymes *ADARB1* and *ADARB2* are associated with extreme old age ^11^. While the functional consequences of these SNPs have not been explored, this association provides further evidence for the importance of A-to-I editing in aging. Our study goes one step beyond previous work focusing solely on ADAR enzymes by demonstrating an age-dependent increase in target RNA editing genome-wide. With this information, it is also important to consider the effects of A-to-I editing on individual target mRNAs. We identified dozens of genes with edits in protein coding sequence that may alter protein function or the speed of translation ^43,44^. Many more inosine sites were detected within non-coding regions of mRNA, where their function is not as well understood, but may affect secondary structure, microRNA targeting, or splicing ^46^.

Many edited nucleotides reside within genes that promote or antagonize longevity. It is therefore possible that loss of A-to-I editing at specific sites could explain the lifespan phenotypes of *adr-1* and *adr-2* loss of function mutants.

### Detection and age-dependent increase of pseudouridine in mRNA

This study is the first to identify and validate pseudouridine edits in *C. elegans* mRNA. Identification of this modification was made possible with Nanopore DRS, highlighting one of the key advantages of this sequencing method. This exciting discovery suggests a previously unknown mode of post-transcriptional regulation in this broadly used model organism and paves the way for future studies investigating the functional roles of Ψ. The age-dependent increase in Ψ and presence of this modification in many aging genes are also intriguing. We identified multiple putative pseudouridine synthase genes in *C. elegans* based on homology to human, which would be exciting targets for future studies investigating the functional roles of Ψ in mRNA and the effect of this modification within specific aging genes. Due to its difficulty to detect and lack of known reader proteins, the role of Ψ in mRNA currently remains elusive.

There is some evidence that Ψ synthase enzymes may recognize structural motifs ^63^ and that Ψ stabilizes RNA-RNA interactions and affects translation ^64^. Ψ in coding regions was shown to alter mRNA translation rate and promote low level synthesis of multiple peptides from a single mRNA in human cells ^65^. As the majority of Ψ sites found in our study are in protein coding regions of mRNA, these modifications could affect protein production for the several hundred genes identified. While the functional consequence of Ψ in mRNA is not well understood, aging *C. elegans* provides a strong model for future studies on the biological relevance of this modification.

### Annotating new features of genes that promote and antagonize longevity

While previous studies have explored changes to the transcriptome that coincide with aging, our datasets and analyses provide an in-depth characterization that goes beyond standard differential gene expression comparisons. Our approach with eight adult collection time points and paired Nanopore DRS and Illumina RNA-seq allowed us to ask new questions with high resolution, leading to identification of novel isoforms, 3’UTRs, and modified RNA nucleotides. The process of aging is multifaceted and complex. Maintenance of normal lifespan is regulated through many pathways important for normal cellular processes, including metabolism, autophagy, apoptosis, cell proliferation, and protein synthesis ^66^. Perturbation of individual genes that control these processes, therefore, can lead to a shortened or extended lifespan. As alternative splicing and RNA modifications can alter transcript abundance, translational efficiency, and function of the encoded protein, this work serves as a valuable resource for advancing a deeper understanding of aging.

### Limitations of the study

This study remains limited by our inability to rely fully on Nanopore DRS data for all the analyses conducted. As aforementioned, the read truncations, lower depth of sequencing, and higher error rates forced us to rely on short reads for some of our analyses. With our approach using two different sequencing methods, however, we were able to overcome many of the challenges posed by Nanopore DRS and make use of this exciting new technology. Because of the newness of this technology, we were careful to use stringent filtering criteria and conservative cutoffs for definition of novel isoforms and pseudouridine. Indeed, we were able to validate our computational predictions experimentally for examined genes. There are likely many more isoforms and sites of interest to be uncovered in our publicly available data.

Recent studies have demonstrated that investigating aging in whole animals may limit our understanding of how gene expression changes relate to functional decline. New technologies have enabled tissue-specific or single-cell RNA-seq in aging *C. elegans* and have shown that the transcriptomes of different tissues or cell types exhibit distinct changes in aging ^67,68^. Therefore, there are likely trends that we miss due to the relatively lower resolution of whole animals. We look forward to future iterations of long-read sequencing that require lower inputs and allow for adaptations of exciting new technologies.

## Acknowledgements

We thank members of the Pasquinelli Lab, H. Cook-Andersen, and E. Yeo for helpful discussions and critical reading of the manuscript. Support for this study was provided by a UCSD Cellular and Molecular Genetics Training Program institutional grant from the National Institute of General Medical Sciences (T32 GM007240) (E.C.S.). This work was funded by grants from the NIH through NIGMS (GM127012) and NIA (AG056562) and the Hevolution Foundation (HF-GRO-23-1199180-41) to A.E.P. Some data were generated at the UC San Diego IGM Genomics Center utilizing an Illumina NovaSeq 6000 that was purchased with funding from a National Institutes of Health SIG grant (#S10 OD026929). This work used Expanse CPU at SDSC through allocation MCB190148 from the Advanced Cyberinfrastructure Coordination Ecosystem: Services & Support (ACCESS) program, which is supported by National Science Foundation grants #2138259, #2138286, #2138307, #2137603, and #2138296.

## Author Contributions

Conceptualization: E.C.S., I.A.N., A.E.P.

Formal analysis: E.C.S.

Funding acquisition: A.E.P.

Investigation: E.C.S., I.A.N.

Methodology: E.C.S., I.A.N.

Supervision: A.E.P.

Validation: E.C.S.

Visualization: E.C.S.

Writing – original draft: E.C.S.

Writing – review & editing: E.C.S. and A.E.P.

## Supplemental information

**Document S1.** Figures S1-S7

**Table S1.** Illumina RNA-seq DESeq2 results from days 2, 3, 4, 5, 7, 10, and 15 vs. day 1.

**Table S2.** Nanopore DRS DESeq2 results using unfiltered reads from days 2, 3, 4, 5, 7, 10, and 15 vs. day 1.

**Table S3.** Number of Nanopore DRS reads per replicate remaining after each filtering step (mapping, transcription start site filtering, and poly(A) filtering).

**Table S4.** All detected Nanopore DRS full-length isoforms with columns 1-12 in BED format and the number of filtered reads for each isoform per replicate. Novel isoforms not in the WS279 WormBase annotation are labeled as productive if they span in frame annotated start and stop codons or unproductive if they do not span in frame annotated start and stop codons.

**Table S5.** 3’UTR coordinates of Nanopore DRS full-length isoforms with columns 1-12 in BED format. 3’UTR sequences are compared to the WS279 WormBase annotation.

**Table S6.** Alternative exon inclusion events with a significant (*P*<0.05) change in relative abundance (PSI) in days 2, 3, 4, 5, 7, 10, or 15 compared to day 1.

**Table S7.** Percent of reads mapping to introns or intergenic regions and percent of splice junction reads that do not match WS279 WormBase annotations in young (days 1, 2, 3) and old (days 7, 10, 15) animals.

**Table S8.** Median, mean, minimum, and maximum poly(A) tail lengths for isoforms with full-length support.

**Table S9.** High confidence A-to-G edit site coordinates in BED format, features that overlap with A-to-G sites, and predicted codon changes resulting from editing.

**Table S10.** High confidence pseudouridine edit site coordinates in BED format and overlapping features.

**Table S11.** Annotated aging genes with novel features from this study indicated.

## STAR Methods

### *C. elegans* strains and collections

*C. elegans* N2 animals were grown under standard laboratory conditions on NGM plates seeded with OP50 *E. coli* ^69^. Synchronization with hypochlorite treatment was performed and embryos were plated and grown at 20°C. Collections were performed at days 1-5, 7, 10, and 15 starting at day 1 of adulthood, 96 h after plating.

### RNA extraction and sequencing

Isolation of total RNA was performed using a standard Trizol protocol. Total RNA was treated with proteinase K (NEB) to increase purity. 12 µg of total RNA were prepared for sequencing using the Direct RNA sequencing kit (cat# SQK-RNA002) and was sequenced on MinION (ONT). The same RNA was also prepared with the Illumina Stranded TruSeq RNA library prep kit. Prior to library preparations, ribosomal RNA was removed using RiboZero Gold (Illumina). cDNA libraries were sequenced on Illumina’s NovaSeq 6000 (100 bp paired-end reads).

### Illumina RNA-seq mapping and read counting

Illumina RNA sequencing reads were aligned to the Wormbase WS279 reference genome using STAR version 2.7.9a ^70^ with parameters --runThreadN 24 --readFilesCommand zcat -- outSAMtype BAM SortedByCoordinate --outFileNamePrefix. Counting was performed with featureCounts ^71^ with parameters -p -s 2. Read normalization and differential gene expression analyses were conducted with DESeq2 ^72^, comparing all time points to day 1. Genes with a Benjamini-Hochberg adjusted *P*-value ≤0.05 were considered differentially expressed.

### Nanopore DRS mapping and filtering

Nanopore RNA sequencing reads were mapped to the Wormbase WS279 reference genome using minimap2 version 2.22-r1101 ^73^ with parameters -a -x splice -uf -k14. To assign unfiltered reads to genes and perform precursory differential expression analysis, featureCounts ^71^ was used with parameters --fracOverlap 0.8 –primary -R CORE -L. Read normalization and differential gene expression analyses were conducted with DESeq2, comparing all time points to day 1. Genes with a Benjamini-Hochberg adjusted *P*-value ≤0.05 were considered differentially expressed. To perform read filtering for full-length isoform identification, primary reads were filtered using samtools version 1.15.1 ^74^ with parameters -F0x100 -F0x900. The TSS_filter.py script ^14^ was used to filter full length reads from primary alignments. Then, filtered reads with a “PASS” flag from Nanopolish poly(A) tail length estimation were used for downstream analyses.

### Splice isoform and 3’UTR analysis

Assigning reads to isoforms and splicing analysis was conducted using FLAIR ^23^, with WS279 genome annotations. Splice junctions from Illumina RNA-seq were obtained using the junctions_from_sam.py script. Nanopore splice junctions were corrected with the flair.py correct module and collapsed using the flair.py collapse module with the -s 0.10 parameter, then isoforms were filtered for a minimum of 20 total assigned reads. The predictProductivity module was used and 3’UTR coordinates were defined from the stop codon to the 3’ end of productive transcripts. 3’UTRs whose 3’ ends did not fall with 10 bp of WS279 WormBase 3’UTR annotations were considered novel. For identification of fusion isoforms, all full-length novel isoforms containing intron(s) longer than the median gene length in *C. elegans* (1,956 bases) and that overlap with exons from two or more annotated WormBase WS279 genes were further examined.

### Long gene fusion analysis

Unfiltered Nanopore DRS alignments were used to run LongGF with default parameters ^35^. Predicted gene fusions with ≥20 supporting reads were kept.

### RT-PCR for novel isoforms

Primers were designed to flank novel splicing events. Reverse transcription of day 1 RNA was performed with SuperScript™III Reverse Transcriptase (Thermo Fisher) and PCR was performed using GoTaq® G2 DNA Polymerase (Promega). PCR products were visualized by agarose gel electrophoresis. Genomic DNA was isolated using a standard Trizol protocol.

#### Primer sequences

*gop-1* Forward: GTAGAAGCGACCGAGGAAGATGA

*gop-1* Reverse: CAGCAGATTGAAATCGAGAACGAGA

*nstp-1* Forward: TCACAGCGACATATGGTCTTATATTT

*nstp-1* Reverse: GCACTGATTTGTTTTAAACTGTACTGA

*aak-1* Forward: AGCAATGGAAGATTTTTGGGAGAT

*aak-1* Reverse: AGATTCCATGTCAATTTCTTTTGCTC

### *lat-1:enol-1* Forward: CCATCTCACACACAATATGTGATGGT

*lat-1:enol-1* Reverse: TGAGAACCCCCTTTCCGAGG

E02D9.1:*mek-5* Forward: ATTTACATTTTGGGACGTGGATCC

E02D9.1:*mek-5* Reverse: ACTGCTCACTTCACAACCATGAT

C18H9.6:*clec-173* Forward: GTAACTCTCGGCCAGGGATG

C18H9.6:*clec-173* Reverse: TCAATCTTTGGGTCAGTCAGAACC

### Alternative splicing analysis

A GTF file of all transcripts identified by Nanopore DRS was created and SUPPA2 ^37^ was used to generate an alternative splicing events file and perform PSI calculations with Illumina RNA-seq data.

### Assignment of reads to introns and intergenic regions

Stringent intronic genomic coordinates were derived from the Wormbase WS279 annotation by pulling regions that lie within transcripts between annotated exons then removing regions that overlap with any annotated exon. Stringent intergenic genomic coordinates were derived from the WS279 annotation by pulling regions that lie between annotated transcripts and removing regions that overlap with any annotated exon. A SAF file was generated for each feature type and Illumina RNA sequencing STAR alignments were assigned using featureCounts with parameters -p -s 2 -T 8 -F SAF --fracOverlap 0.5.

### Splice junction fidelity analysis

A database of intronic genomic coordinates was created from the Wormbase WS279 annotation by pulling regions that lie within transcripts between annotated exons. High confidence collapsed splice junction coordinates from STAR mapping (SJ.out.tab files) were compared to the intronic database. Uniquely mapping reads that spanned annotated or unannotated splice junctions were counted and a percentage of unannotated splice junctions was derived by dividing unannotated splice junction counts by the total number of uniquely mapping reads spanning splice junctions.

### Poly(A) tail length analysis

Nanopolish version 0.13.3 ^75^ was used to index fast5 and fastq files and perform poly(A) tail length calling. Filtered reads with “PASS” tags were used for analysis of poly(A) tail lengths with read assignments from FLAIR.

### Inosine detection genome-wide

The SAILOR pipeline from the FLARE package (https://github.com/YeoLab/FLARE) was used to identify A-to-G edits genome-wide. Sites with a proportion of edited reads ≥0.05 and a quality score ≥0.75 in three independent biological replicates were used for downstream analyses.

### Pseudouridine detection genome-wide

NanoPsu ^48^ was used to identify putative pseudouridine sites in unfiltered Nanopore DRS reads. Sites with a probability score ≥0.90 in three independent biological replicates were used for downstream analyses.

### Validation of pseudouridine sites

Bisulfite/sulfite treatment was performed following a published protocol with minimal modifications ^52^. Briefly, 1 µg of DNase treated RNA pooled from adult days 1, 2, and 3 collections was sulfite/bisulfite treated, then reverse transcribed with Maxima H Minus (Thermo Fisher Scientific, EP0753) with random primers (New England Biolabs, S1230S) according to manufacturer’s instructions, alongside an untreated control. PCR with gene specific primers flanking putative Ψ sites was performed with GoTaq® G2 DNA Polymerase (Promega, M7841). PCR products were recovered and cloned into TOPO-TA cloning vectors and transformed into *E. coli* competent cells. Sanger sequencing was performed for at least 5 individual clones.

#### Primer sequences

*rnh-2* Forward: CAATTTCGGGAATGGTATTCCGT

*rnh-2* Reverse: TCGAATCAAAGTGATCGCAGC

*rrn-1.1* Forward: GACTACCATGGTTGTTACGGGTAA

*rrn-1.1* Reverse: GGCATCGTTTACGGTCAGAACTAG

## Data and code availability

- All datasets have been deposited at Sequence Read Archive (SRA) and will be freely available at the time of publication.
- All original code has been deposited at Zenodo under: https://zenodo.org/doi/10.5281/zenodo.11672730
- Any additional information required to reanalyze data from this study is available upon request from the lead contact, Amy Pasquinelli

## Materials availability

This study did not generate new unique reagents.

