## Supplemental Figures and Tables for "Full-length direct RNA sequencing reveals extensive remodeling of RNA expression, processing and modification in aging *Caenorhabditis elegans*": Schiksnis_Cell_Reports_Document_S1.pdf

### Supplementary Figures

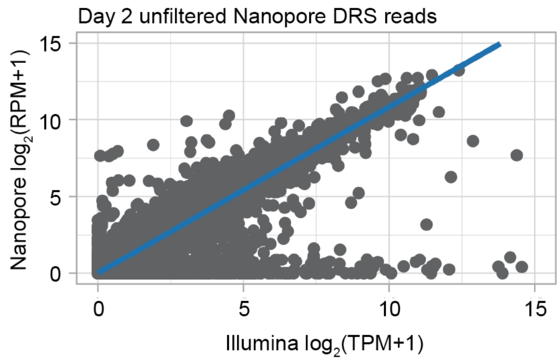


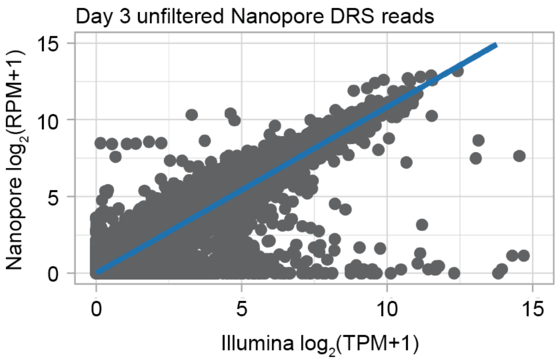


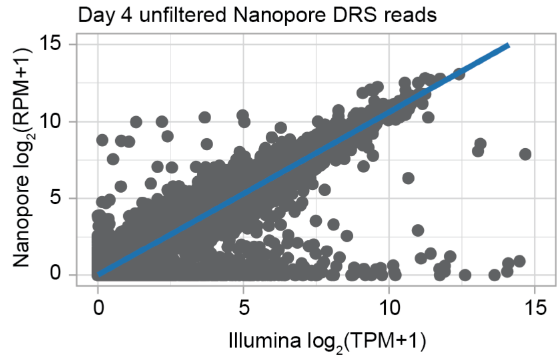


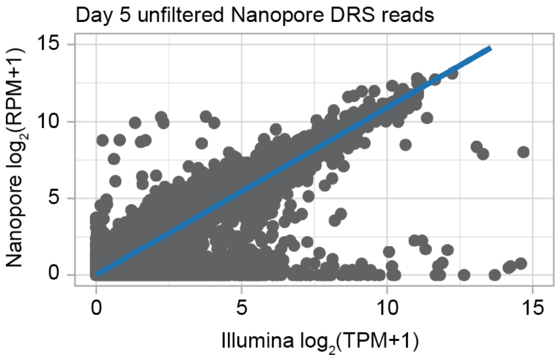


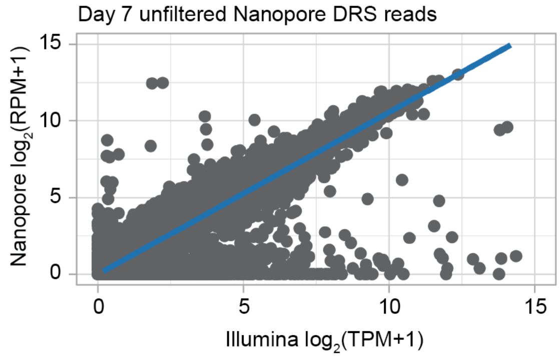


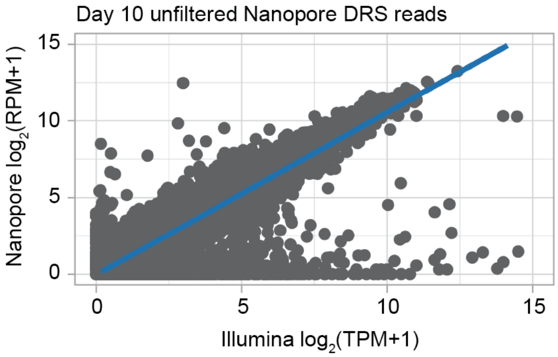


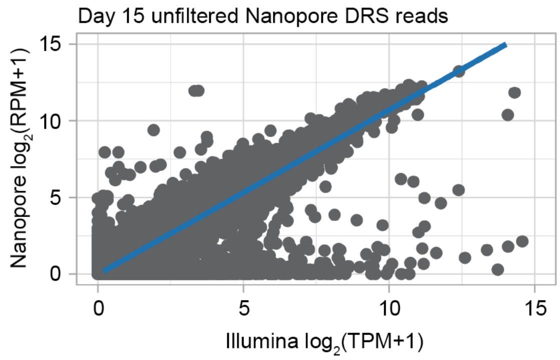

**Figure S1, related to Figure 1. Correlations between Nanopore DRS and Illumina RNA-seq normalized gene expression.** Scatterplots comparing mean normalized expression of individual genes across three independent biological replicates between Nanopore DRS ($log_2$ TPM + 1) and Illumina RNA-seq ($log_2$ RPM + 1) at each time point.

A B

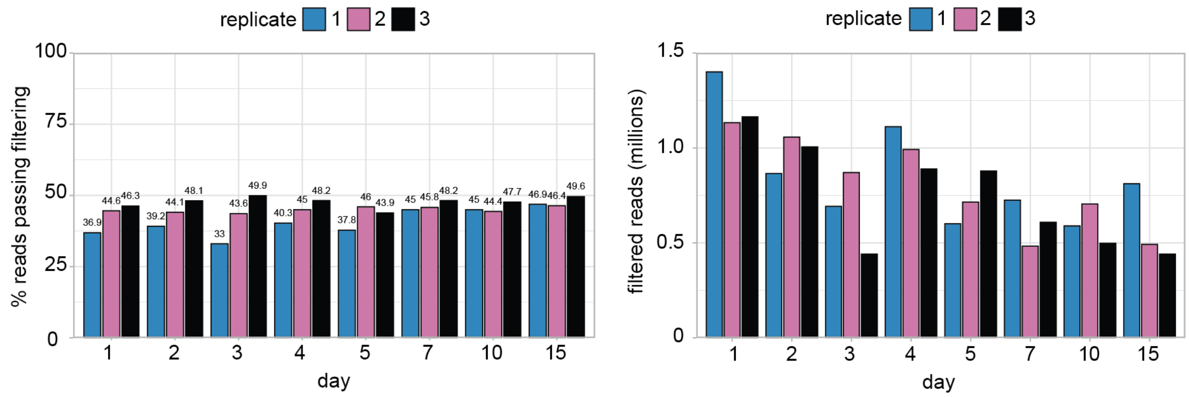


C

| Day | Nanopore (full-length only) and Illumina correlation ($R^2$) |
|---|---|
| 1 | 0.51 |
| 2 | 0.53 |
| 3 | 0.52 |
| 4 | 0.54 |
| 5 | 0.54 |
| 7 | 0.50 |
| 10 | 0.48 |
| 15 | 0.48 |

D

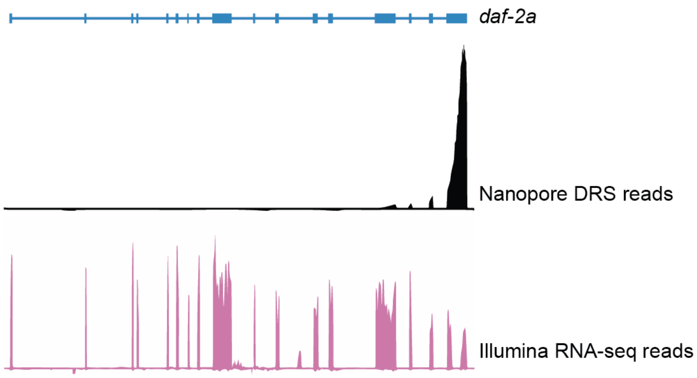


**Figure S2, related to Figure 2. Read filtering for Nanopore DRS. A.** Percent of reads that remain after Nanopore filtering pipeline at each time point. **B.** Number of reads that remain after Nanopore filtering pipeline at each time point. **C.** Table showing the R-squared value for mean normalized expression for filtered Nanopore DRS ($\log_2$ TPM + 1) and Illumina RNA-seq ($\log_2$ RPM + 1) of individual genes across three independent biological replicates at each time point. **D.** Browser track image of *daf-2a* showing unfiltered Nanopore DRS and Illumina RNA-seq read density.

**A**

| Gene fusion | Total counts | Break 1 | Break 2 |
|---|---|---|---|
| C18H9.6:*clec-173* | 728 | II:6701465 (-) | IV:3266420 (-) |
| T28F3.8:*clec-84* | 293 | IV:17307321 (-) | IV:2845291 (+) |
| Y51H4A.24:*clec-84* | 62 | IV:16739384 (-) | IV:2845291 (+) |
| *eri-6*:*eri-7* | 57 | I:4458645 (+) | I:4456339 (-) |
| *clec-84*:K08D10.14 | 66 | IV:2845291 (+) | I:4456339 (-) |

**B**

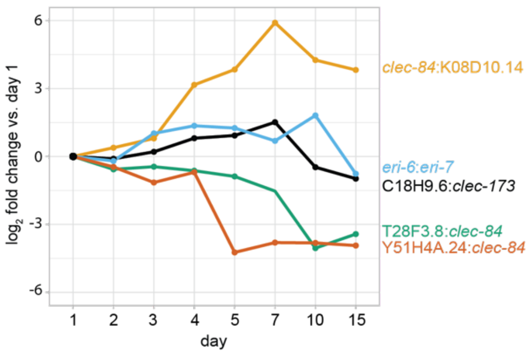


**C**

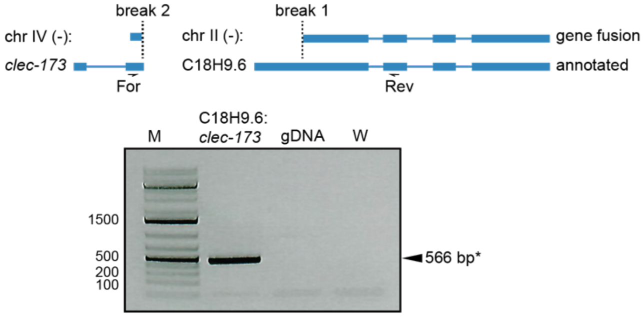


**Figure S3, related to Figure 2. Detection of gene fusion isoforms at adult time points. A.** Table of gene fusions detected with LongGF. Break points indicate the genomic coordinates where two genes are joined. **B.** Line plot showing the relative expression normalized to day 1 of LongGF fusion genes. **C.** RT-PCR validation of the C18H9.6:*clec-173* LongGF fusion isoform. Agarose gel electrophoresis (*bottom*) with expected fragment size denoted, including genomic DNA and water controls. Schematic representations of gene fusion (*top*) showing break points and forward and reverse PCR primers used for amplification of cDNA.

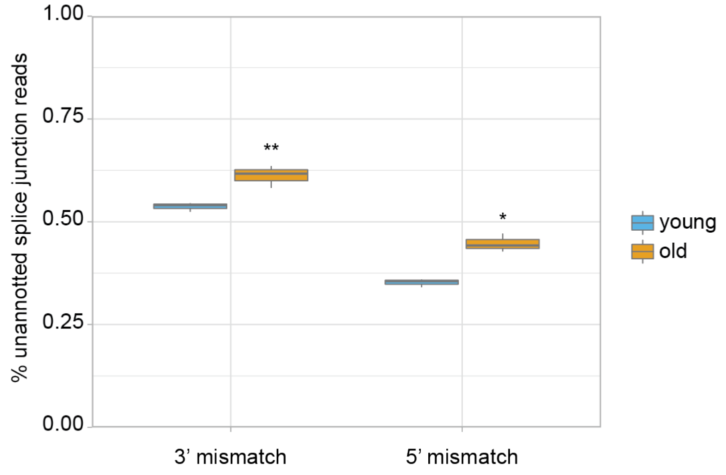


**Figure S4, related to Figure 4. Percent of splice junction reads using unannotated 5’ or 3’ splice sites.** Boxplot showing the percent of splice junction reads that do not match WS279 annotated 5’ splice sites (**$P$=0.004, *t*-test, two-sided after probit transformation) or 3’ splice sites (*$P$=0.022, *t*-test, two-sided after probit transformation).

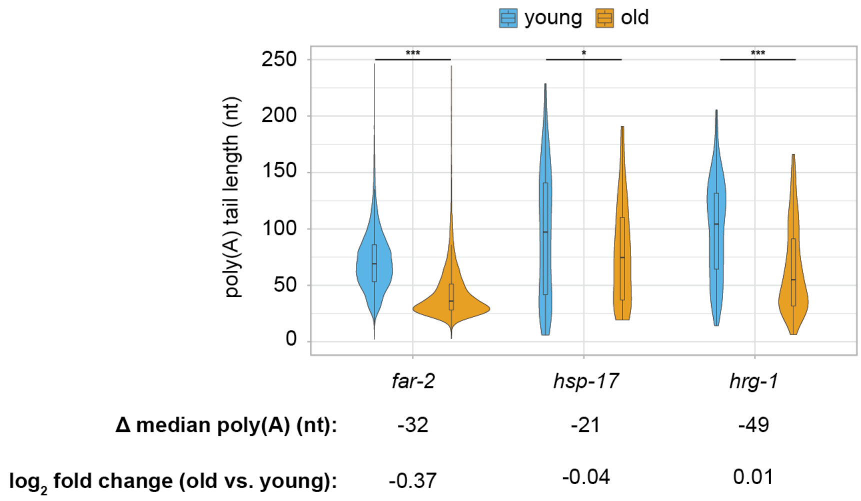


**Figure S5, related to Figure 4. Genes with large changes to poly(A) tail lengths in aging.** Violin plot with inlaid boxplot of poly(A) tail length distribution for reads assigned to indicated genes. Changes to media poly(A) tail lengths and expression in old compared to young animals are indicated. ****P*<0.001, **P*<0.05 (Kolmogorov-Smirnov test) between young and old animals.

**A**

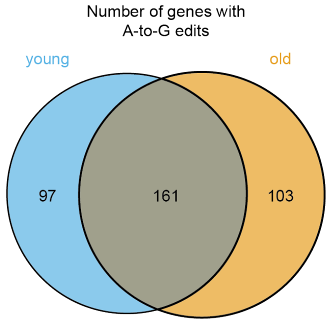


**B**

| Amino acid group | Amino acids |
|---|---|
| Positively charged | R,H,K |
| Negatively charged | D,E |
| Polar | S,T,N,Q |
| Hydrophobic | A,V,I,L,M,F,Y,W |
| Special | C,G,P |

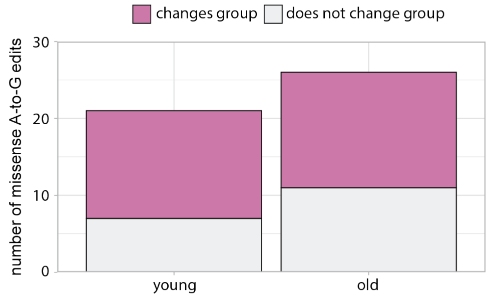


**Figure S6, related to Figure 5. Examination of A-to-I editing in aging. A.** Venn diagram with the number of genes with A-to-G edits in young and old animals. **B.** Table with amino acid group characterizations based on chemical properties (*left*) and stacked bar plot showing the number of predicted missense A-to-G edits that result in a change to the encoded amino acid group (*right*).

**A**

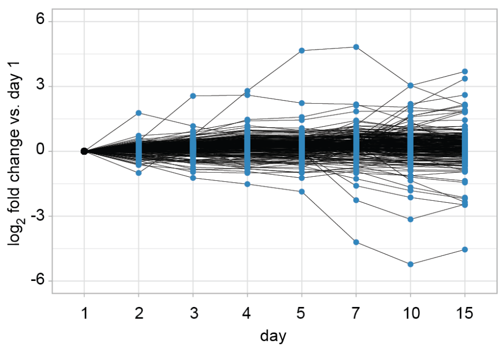


**B**

| Gene | Human homolog | Target(s) of human homolog |
|---|---|---|
| *pus-1* | Pus1 | tRNA |
| *tag-124* | Pus3 | tRNA, mRNA |
| Y43B11AR.3 | TruB2 | tRNA, mt-mRNA |
| K07E8.7 | RpusD2 | mRNA |
| ZK287.7 | RpusD3, RPusD4 | mt-rRNA, tRNA |
| B0024.11 | Pus7 | tRNA, mRNA |
| Y48C3A.20 | Pus10 | tRNA, rRNA, snRNA |

**C**

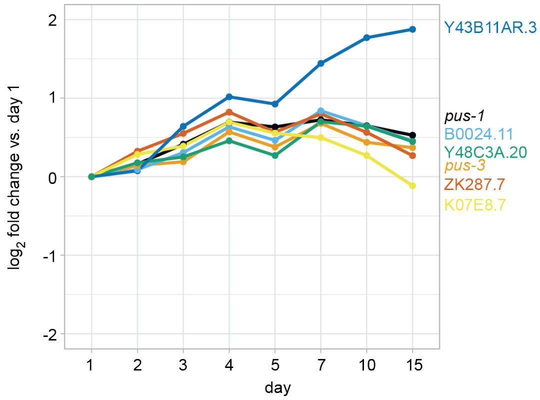


**Figure S7, related to Figure 6. Dynamic expression of genes with pseudouridine and pseudouridine synthase enzymes. A.** Line plot showing the relative expression normalized to day 1 of genes with pseudouridine in old animals but not young. Each blue dot represents an individual gene. **B.** Table of all predicted *C. elegans* homologs of human pseudouridine synthase enzymes and their RNA targets in human. **C.** Line plot showing the relative expression normalized to day 1 of predicted pseudouridine synthase genes.
